# A case for slow-wave morphology as a functional biomarker of gastric disease

**DOI:** 10.1101/2025.05.21.655419

**Authors:** Jarrah M. Dowrick, Jonathan C. Erickson, Peng Du, Timothy R. Angeli-Gordon

## Abstract

**Aim:** Underlying bioelectrical slow waves are critical for regulating gastric motility, and abnormal spatiotemporal slow-wave ‘dysrhythmias’ are associated with a range of gastrointestinal disorders. However, the definition and role of the morphology of gastric slow-wave signals have remained limited. This study aimed to investigate the potential of gastric slow-wave morphology as an actionable biomarker.

**Methods:** Data were repurposed from a study where, following ethical approval, a control cohort (*n*=9) and a pathological cohort of patients with chronic unexplained nausea and vomiting (CUNV; *n*=8) underwent intra-operative high-resolution serosal electrical mapping (96-256 electrodes, 4.0-5.2 mm spacing). Slow waves were identified using validated software, and spatiotemporally averaged waveforms were compared between cohorts. These waveforms were replicated in a computational model of gastric slow-wave propagation to explore potential functional implications.

**Results:** The slow-wave morphology of the CUNV cohort exhibited a more gradual recovery stroke compared to controls, which manifested as an increase in the normalized recovery stroke area [0.206 (95% CI: 0.169–0.247) vs. 0.134 (95% CI: 0.106–0.166); *p*=0.011]. Computational modeling showed that these morphological differences could drive spatial slow-wave dysrhythmias. Considering the evident functional importance of gastric slow-wave morphology, we highlighted the three typical morphological features: 1) rapid, brief upstroke, 2) downstroke, and 3) biphasic recovery stroke.

**Conclusion:** Altogether, this study presents a case for gastric slow-wave morphology as a biomarker of gastric health and disease and lays a foundation for the standardization of future slow-wave morphology research.

**PRACTITIONER POINTS:**

- Abnormal spatial slow-wave propagation is associated with a range of gastrointestinal motility disorders, but morphology has had limited consideration.
- Slow-wave morphology differs between cohorts of healthy controls and patients with chronic unexplained nausea and vomiting.
- Multiscale mathematical modeling indicates that a disruption to the slow-wave recovery stroke may contribute to spatial disorganization.

## INTRODUCTION

The frequency and spatial propagation of gastric slow waves, electrical events generated and propagated via an interconnected network of interstitial cells of Cajal (ICC), are proven valuable diagnostic biomarkers for pathologies like gastroparesis (1–3) and chronic nausea and vomiting (4–6). Consequently, much effort has been invested in developing devices and techniques to enhance the accessibility and quality of gastric electrical measurements, including intra-operative (7,8), endoscopic (9–13), and body-surface approaches (14–16). Even with the evolution of slow-wave recording technology and proliferation of data on bioelectrical slow-wave dysrhythmias underlying motility disorders (17,18), the definition and functional implications of gastric slow-wave morphology remain limited.

In contrast, cardiology has successfully integrated electrophysiological measurement of the electrocardiogram (ECG) into routine clinical practice (19,20), providing inspiration for how gastric electrophysiology might evolve. Curiously, both the ECG and gastric electrophysiology can trace their origin to the early 20^th^ century, when Prof. Walter Alvarez recorded the first documented gastric slow wave (21) using a galvanometer setup similar to the one used by Einthoven to capture the first ECG only decades earlier (22,23). Despite the parallels in their origin, there has been a clear divergence in the maturation of the ECG and gastric slow waves.

Various aspects of slow-wave morphology, including amplitude, downstroke width, and recovery duration (7,24,25), have been used to quantitatively characterize gastric activity in research applications, but they have seen limited clinical uptake. Comparatively, the signal morphology of the electrocardiogram (ECG) in the heart has been comprehensively defined (*i.e.*, the PQRST complex (26)) and incorporated into routine clinical practice (19,20). Waveform characteristics and variations of the ECG serve as a valuable diagnostic tool in cardiology, where the ECG is recommended as a diagnostic biomarker of ventricular arrhythmia (27), ischemia (28), hypertrophy (29), and many other disorders (20), all associated with defined variations of the ECG waveform, like the ST segment or QT interval (30). Defining typical gastric slow-wave morphology could enable it to serve as an actionable biomarker of gastric function, holding clinical relevance in diagnosing, monitoring, and treating gastric motility disorders, as the ECG does in the heart.

In this study, we performed a cohort-wise comparison of extracellular gastric slow-wave signal morphology using data from previously reported cohorts of controls and patients with chronic unexplained nausea and vomiting (CUNV). We subsequently investigated the functional implications of changes in slow-wave morphology using *in silico* simulations. We hypothesized that: i) gastric slow waves have identifiable morphological features, ii) morphological differences exist in healthy versus diseased cohorts, and iii) local changes in slow-wave morphology can influence spatial bioelectrical propagation patterns.

## RESULTS

### Study population

The characteristics of each cohort have been reported previously (4). Briefly, the CUNV cohort consisted of eight patients (7 female and 1 male), and the control cohort consisted of nine subjects (5 female and 4 male). The total number of marked waves was 7,595 for the CUNV cohort and 7,761 for the healthy controls.

### Slow-wave morphology in health and CUNV

Morphology varied within and between cohorts (Figure 1), though the average waveform of all subjects could be summarized as consisting of: i) an initial upstroke, ii) a brief, rapid downstroke, and iii) a slower recovery stroke. The recovery stroke contained much of the variability, particularly among control subjects. Some controls presented with a monophasic recovery period (Control 1, 4, and 7; Figure 1), while the majority exhibited a biphasic morphology, with an initial rapid recovery, followed by a more gradual convex trajectory (Control 5; Figure 1). This recovery period was more consistent among CUNV patients, with most exhibiting a minimal rapid phase and a more exaggerated convex return to baseline.

**Figure 1:**
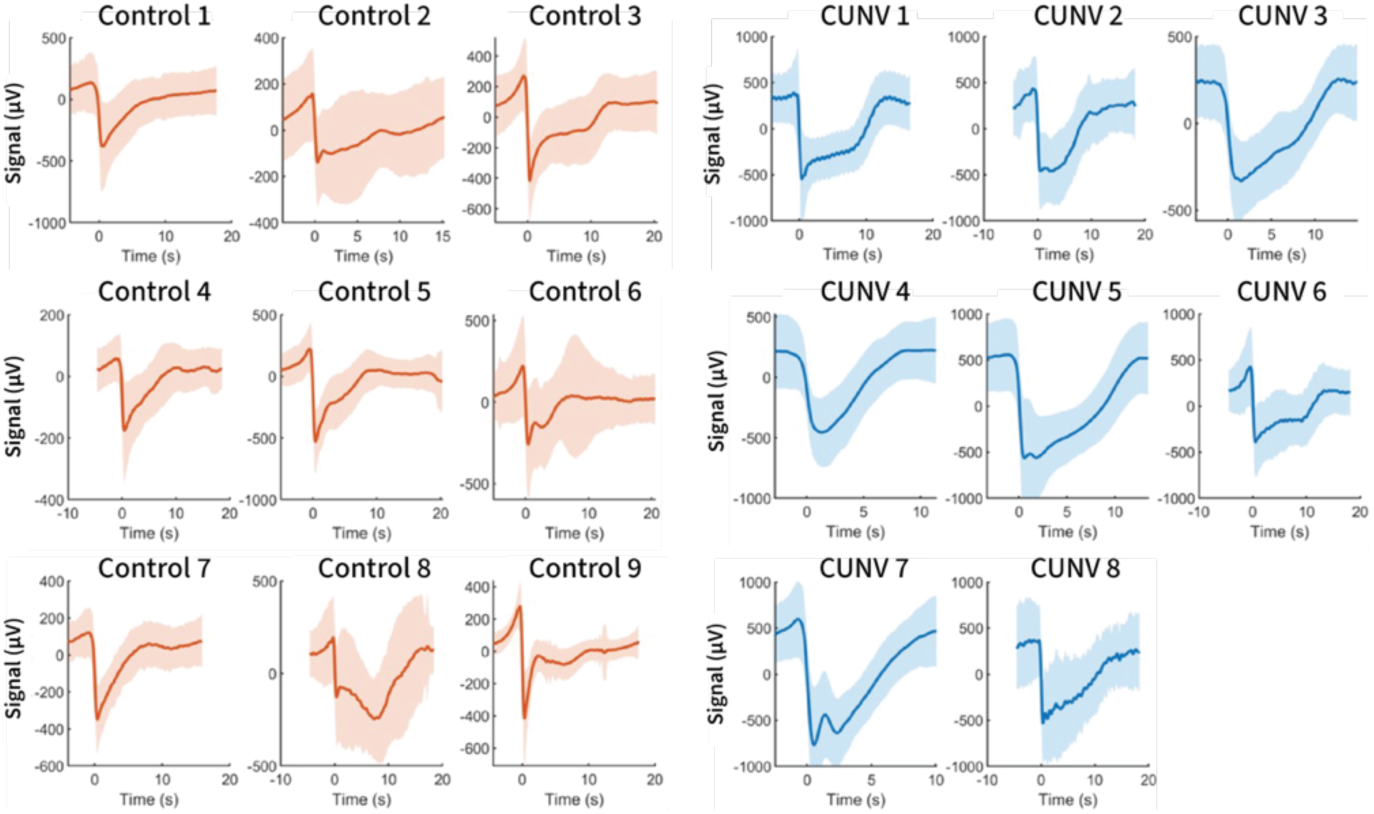
Subject-wise average slow-wave morphology. Each panel contains the array-wide averaged slow-wave signal from a single subject (thick line). The shaded region denotes the standard deviation of the signals. Average slow-wave signals from chronic unexplained nausea and vomiting (CUNV) subjects are arranged on the right and colored blue, and those from control subjects are arranged on the left and colored orange.

When considering the cohort-wise median signals (Figure 2), the differences between cohorts became increasingly apparent. The upstroke and downstroke of the waves were largely similar, but the recovery stroke clearly differed. The average control slow-wave signal had a notable two-stage recovery stroke, starting with a relatively fast upstroke and a secondary, slower, concave return to baseline. While the average CUNV slow-wave signal also exhibited a two-stage recovery period, the initial faster segment was much less pronounced, and the subsequent slower return to baseline was convex.

**Figure 2:**
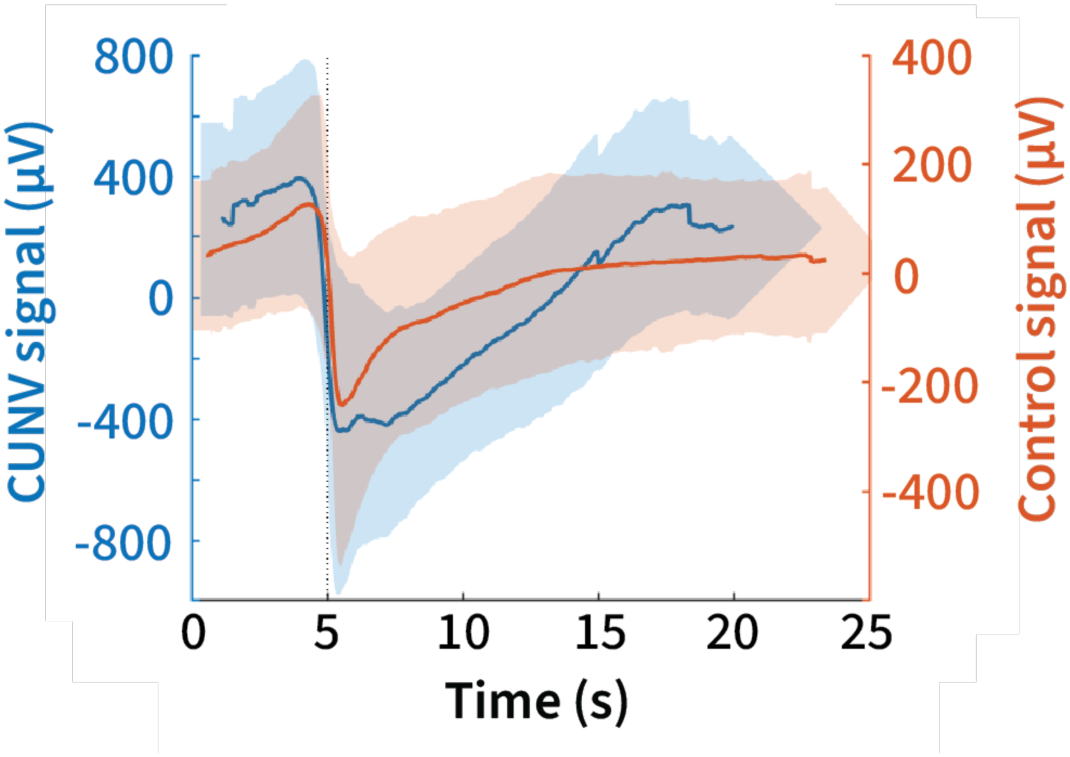
Cohort-wise median slow-wave signal. The median signal for each cohort is described by the thick line. The shaded area surrounding the traces denotes the standard deviation. Different scales and colors are used for each cohort (left/blue – chronic unexplained nausea and vomiting (CUNV), right/orange – control).

In alignment with the cohort-wise median signals, CUNV slow waves had a greater amplitude [671 µV (95% CI: 515–847 µV) vs. 420 µV (95% CI: 306–552 µV); *p*=0.0302; Figure 3A] and shorter downstroke width [0.584 s (95% CI: 0.523–0.645 s) vs. 0.729 s (95% CI: 0.671–0.787 s); *p*=0.0043; Figure 3B]. Normalized RSA was also greater among the CUNV cohort [0.206 (95% CI: 0.169–0.247) vs. 0.134 (95% CI: 0.106–0.166); *p*=0.011; Figure 3C], but there was no difference in the recovery duration [CUNV: 6.74 s (95% CI: 5.77–7.71) vs. control: 5.75 (95% CI: 4.83–6.67 s); *p*=0.17; Figure 3D].

**Figure 3:**
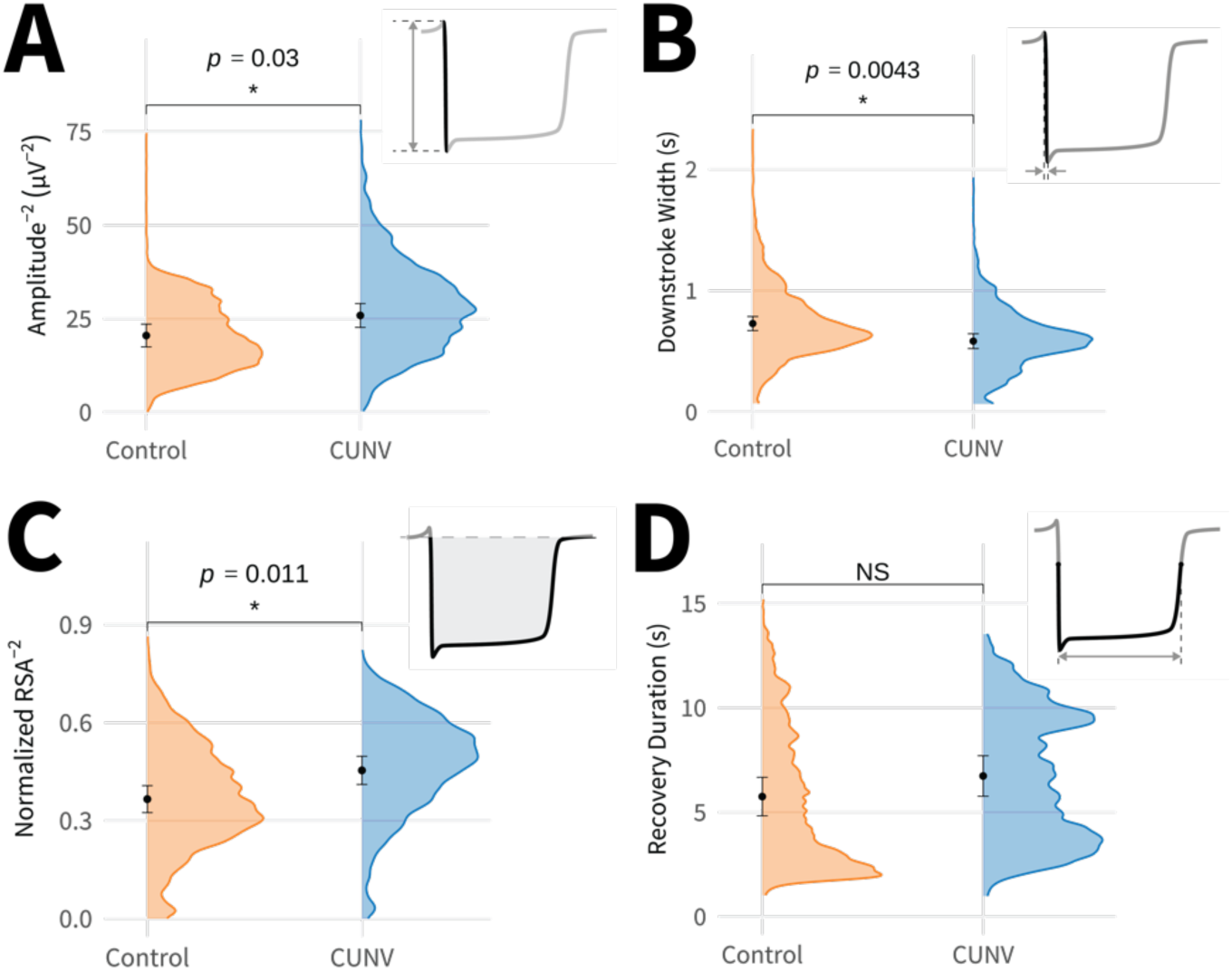
Cohort-wise morphological measure comparison. **(A)** Amplitude, **(B)** downstroke width, **(C)** normalized recovery stroke area (RSA), and **(D)** recovery duration were compared between cohorts. On each panel, the fixed effect (black dot) and 95% confidence intervals (black error bars) are included for each cohort as estimated using the R package *emmeans* (39). The color-coded raw data distribution is also included (orange – control, blue – CUNV). Inlays in the upper right of each panel illustrate the measure being compared. *P*-values were calculated by the R package *lmerTest* (38), and significance was assigned at *p*<0.05. Amplitude and normalized RSA were square root transformed to achieve normally distributed model residuals.

### Multiscale mathematical modeling

Control slow-wave morphology was achieved using the default model parameters, and an approximation of the CUNV slow-wave morphologies was achieved by slowing down the dynamic rate of each slow wave by 64%, which slowed the recovery phase (Figure 4A). When using the *in silico* approximation of the CUNV slow-wave morphology to drive the 1D model, the stable propagation pattern resulting from the default parameter set (Figure 4B) was no longer achieved, and simultaneous antegrade and retrograde wavefronts were present (Figure 4C). Extending the model to 3D affirmed these phenomena (Figures 4D-E), with severe disruptions to normative concentric slow-wave propagation. Animations of the 3D simulation results are included as Supplementary Animation S1 & S2.

**Figure 4:**
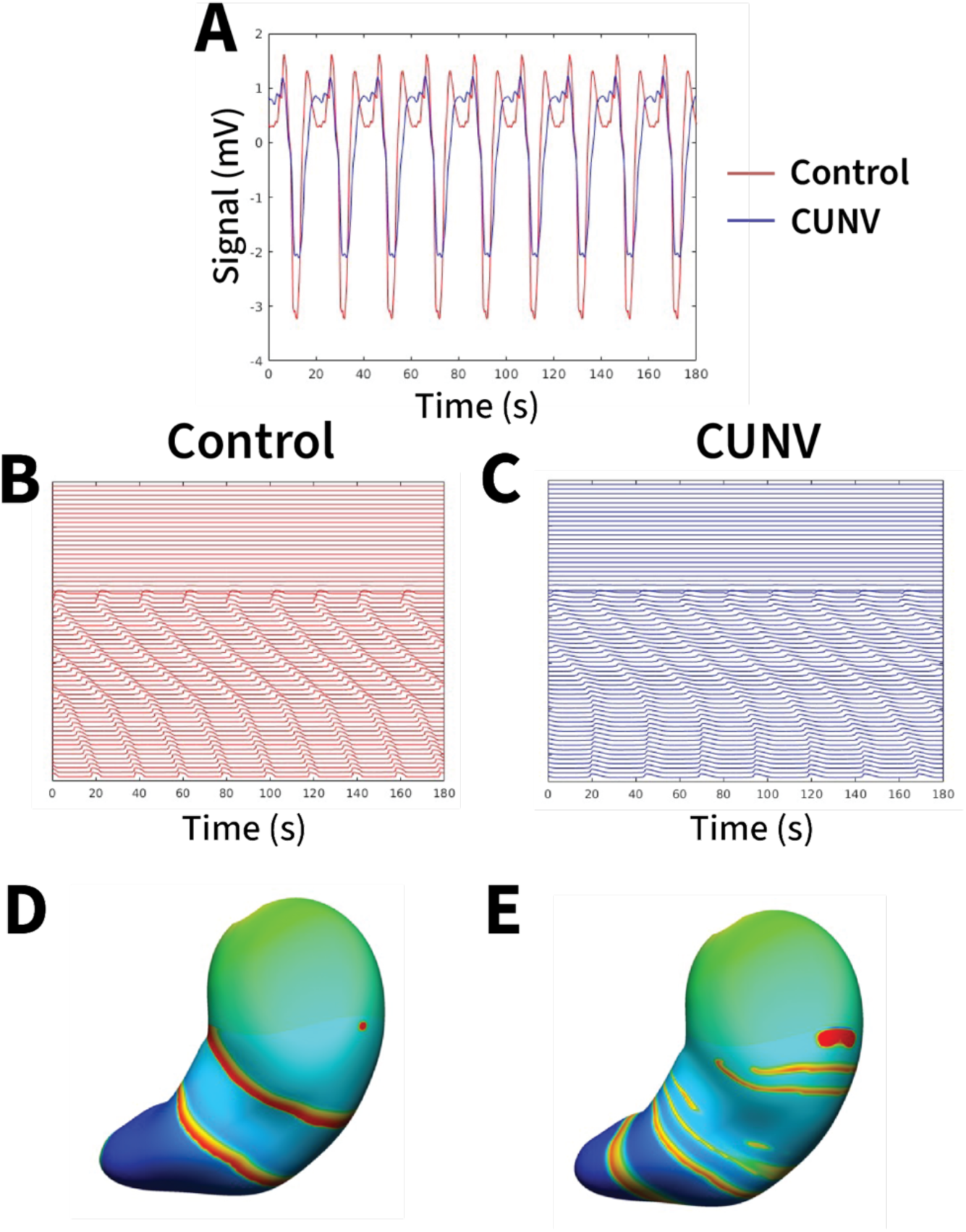
Functional implications of altered slow-wave morphology. **(A)** Extracellular potentials from the midpoint of the 1D model at steady state, approximating the average slow-wave signals from controls (orange) and CUNV patients (blue). **(B)** 1D model steady-state result when using the default (left) and **(C)** CUNV (right) cell model parameters. **(D)** 3D model steady-state result of slow wave propagation pattern when using the default (left) and **(E)** CUNV (right) cell model parameters.

### Standardizing slow-wave morphological feature nomenclature

Given the potential influence of local slow-wave morphology on the spatial propagation of gastric electrical activity, we considered it important to highlight the typical morphological features of extracellular gastric slow waves (Figure 5). The three phases were: i) an initial upstroke, followed by ii) a brief, rapid downstroke, and finally iii) a slower recovery stroke. The recovery stroke was typically biphasic, with an initial rapid recovery phase, followed by a more gradual, secondary return to baseline.

**Figure 5.**
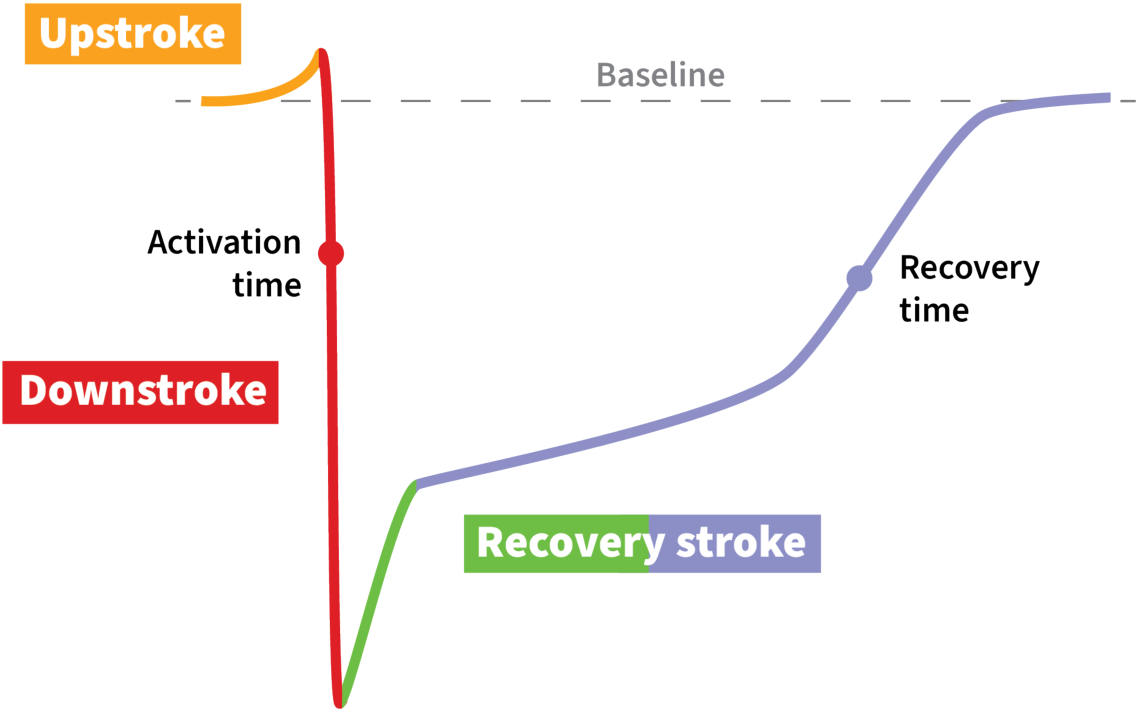
Key features of extracellular gastric slow-wave morphology. Each phase of the slow wave is labeled with different colors, as indicated by the labels: upstroke – yellow, downstroke – red, and recovery stroke – green and purple. The two colors of the recovery stroke reflect the typical biphasic nature of this morphological feature, with a rapid initial recovery (green) and a more gradual secondary return to baseline (purple). Activation time and recovery time are labeled and denoted by red and purple dots, respectively.

## DISCUSSION

This study defined the typical waveform morphology of human extracellular gastric slow waves using *in vivo* HR mapping data collected in cohorts of controls and patients with CUNV. Striking morphological differences in the recovery stroke between these cohorts were captured by a new metric, normalized RSA, along with differences in well-established metrics of amplitude and downstroke width. *In silico* simulations suggested that these morphological differences may contribute to the abnormal slow-wave activity observed in CUNV patients.

Mechanistic links between electrophysiology and motility symptoms have proven evasive despite decades of discoveries associating electrophysiological dysrhythmias with motility disorders, including functional dyspepsia (41–44), gastro-oesophageal reflux (45,46), gastroparesis (1–3), and CUNV (4–6). Gastric Alimetry, with its simultaneous collection of symptoms and body-surface electrical mapping data, has further strengthened the apparent association between electrophysiological dysfunction and gastric disorders (15,16,47). However, the continued limited mechanistic understanding has stunted the development of effective interventions and treatments (48–51). The *in silico* data presented here suggest functional consequences of the recovery stroke. Slowing the recovery stroke disrupted the regular antegrade propagation of electrical events in the 1D and 3D models, effectively representing the spatial dysrhythmias observed in the *in vivo* high-resolution electrical mapping data collected from patients with CUNV (4).

The possible significance of the recovery stroke motivates understanding the underlying, contributory mechanisms. Extracellular electrophysiology reflects not only the local cellular events but also the spatial propagation of electrical activity at a tissue level (8). As such, inter-cohort differences are likely the result of a combination of factors. At the cellular level, the recovery stroke represents the plateau and repolarization phases of intracellular electrical events. Plateau phase duration is primarily influenced by the duration of Ca^2+^ transient clusters (52), with the resultant Ca^2+^ currents ultimately driving the sustained activation of ANO1 Cl^-^ channels (53,54). The Na^+^-Ca^2+^ exchanger (NCX) is also thought to contribute to plateau phase duration, switching into reverse-mode operation under certain conditions, enabling additional Ca^2+^ entry. Disruptions to ICC Na^+^ handling have been shown to influence the amplitude and duration of intracellular slow waves (55). The data of this present study could suggest the presence of a channelopathy, which contributes to the pathophysiology of CUNV, much like the *SCN5A* gene mutation that disrupts the channel Na_v1.5_ and is associated with functional dyspepsia (56).

At the tissue level, extracellular electrical measurements can be considered as the spatial integral of intracellular signals (8). Therefore, tissue characteristics that influence the transmission of electrical activity, such as conductance and capacitance, are critical determinants of extracellular slow-wave morphology. While the *in silico* simulations of this present study exhibited spatial dysrhythmias without changing conductance or capacitance, we cannot disregard their possible contribution, especially given the disrupted ICC network in CUNV patients (4) that would likely influence conductance. Methods like electrical impedance spectroscopy, which are capable of characterizing the electrical properties of tissue, have already been successfully applied to gastric biopsies for cancer (57) and could similarly provide insight into tissue-level disruptions underlying gastric motility disorders.

Downstroke amplitude also differed between cohorts, with the CUNV amplitude almost double that of controls. Initially, this appears to contradict the original data on which this present investigation was built (4), as no difference in amplitude was previously observed. However, the original analysis considered longitudinally propagating slow-wave data only, whereas the current analysis leveraged the entire mapped region. CUNV propagation was typically dysrhythmic (4), and included regions of circumferentially propagating wavefronts. Circumferential propagation occurs with higher velocity (58–60), which is similarly accompanied by higher amplitude in extracellular recordings as more transmembrane current enters the extracellular space with rapid slow-wave propagation (50). Therefore, the observed difference with historical data was anticipated, and the presence of dysrhythmic circumferential propagation in the CUNV cohort may be a key contributor to the morphological differences. As an aside, the ability to resolve between CUNV and control slow-wave morphology without partitioning based on the axis of propagation could be seen as a benefit, reducing the number of electrodes required for insightful chronic monitoring of gastric electrical activity.

Translating such insights from basic science into clinical tools is critical, as gastroenterology is in desperate need of actionable biomarkers (48). Many functional and motility disorders rely on a diagnosis of exclusion, with numerous invasive tests and extended periods of frustration for patients and clinicians alike (61). Gastric slow-wave propagation has already been highlighted as a valuable clinical biomarker (1,4,16), but there has been hesitancy to consider slow-wave morphology for this purpose due to lingering concerns of signal quality (62). To address this, we proposed a new measure, normalized RSA, an integral-based approach. Integral-based measures are robust for quantifying low SNR signals and multi-phasic (‘fractionated’) waveforms, avoiding computational challenges and pitfalls inherent with single-point-based time measures, which often rely on finding maxima or minima in the slow wave signal or its time derivatives (24,32). As such, normalized RSA, particularly averaged over several slow-wave events, may represent a next-generation actionable gastric biomarker.

Activation-recovery interval (here called recovery duration) first drew attention to the association between a dysregulated recovery stroke and spatial slow-wave dysrhythmias (24), representing an important advancement in the morphological analysis of slow-wave morphology. It is defined as the interval between the activation and recovery times, each of which is, in turn, defined using the peak absolute gradient (25,33). However, as demonstrated in this present study, the complex morphology of the recovery stroke corresponds with highly variable estimates of recovery time and, ultimately, in a measure that failed to distinguish between CUNV and control. This result emphasizes the inherent challenges with single-point-based time measures and supports the need for more stable, integral-based measures, such as normalized RSA.

In considering gastric slow-wave morphology as a biomarker, we highlighted the key morphological features and proposed standardized nomenclature to support consistency across the field moving forward. The proposed standardization of slow-wave nomenclature provides a foundation for biomarker investigation and discovery, akin to what is done in the cardiac field with the PQRST complex (27,28). Future morphology studies can build on the observations of an increased downstroke amplitude, smaller initial rapid recovery stroke, and more convex secondary recovery stroke phase in CUNV patients compared to controls.

While this study provides important insights, it is not without limitations. For example, the data were collected using interoperative serosal electrodes in an acute setting. Direct contact recordings from the serosal surface of the stomach are valuable because they give us ground truth slow-wave activity and signal, directly from the source. However, their invasive nature precludes their use as a routine clinical tool, especially as part of a diagnostic screening pipeline. Expanding to establish morphological standards for additional disease cohorts and leveraging emerging methods like implantable monitoring (12,63), minimally invasive endoscopic mapping (9,13), and body-surface gastric mapping (14,15) will help elucidate the scope of slow-wave morphology as a diagnostic biomarker. Second, the array placement may not be consistent between all subjects. As this was a reanalysis of historical data collected from two independent cohorts, we cannot guarantee consistent definitions of anatomy. In screening the recordings, care was taken to only include data collected at or immediately next to the corpus-antrum border. Slow-wave activity is known to exhibit regional variation across the stomach (64–66), so regional standards in health and disease would also be beneficial in the future. Finally, the cohort sizes of eight patients with CUNV and nine healthy controls are relatively small. However, the high-resolution mapping methods yielded over 7,000 individual slow-wave events from each cohort, and significant differences were observed in three of the four morphological measures, reinforcing the consistent inter-cohort differences.

## METHODS

The data reported here have been repurposed from an earlier study (4). Approval was granted by the New Zealand Health and Disabilities Ethics Committee, Auckland District Health Board Research Review Committee, and Institutional Review Board at the University of Mississippi. All patients provided their informed consent.

### Study population

#### Control cohort

Patients undergoing elective upper-abdominal surgery at Auckland City Hospital (New Zealand) were invited to participate as controls. Patients were excluded from this control cohort if they had previously undergone gastric surgery or had a condition or pathology associated with dysrhythmic gastric activity, such as gastroparesis or functional dyspepsia (18). Patients were also excluded if they were taking medications that can affect gastric electrical activity (*e.g.*, opioids).

#### CUNV cohort

Consecutive patients with at least six months of continuous CUNV symptoms undergoing implantation of a gastric electrical stimulation device at the University of Mississippi Medical Center (United States) were invited to participate. All patients had a clinical history, drug history, physical examination, upper endoscopy, and appropriate radiologic and laboratory investigations to exclude other causes of symptoms. Patients with cyclic vomiting syndrome, gastroparesis, malignancy, primary eating disorders, or pregnancy were excluded. As hyperglycemia can induce slow-wave dysrhythmias (31), blood glucose levels were kept within the normal range during the perioperative period.

### Data collection

Following general anesthesia and upper-midline laparotomy, and before organ handling or stimulator placement, high-resolution mapping data were collected for up to 15 min. During this window, a flexible-printed-circuit (FPC) electrode array (96-256 electrodes; 4.0-5.2 mm inter-electrode spacing; 12-59 cm^2^) was placed on the anterior serosal surface of the stomach in the corpus or corpus-antrum border region. Unipolar recordings were acquired from the FPC electrode array at 512 Hz using an ActiveTwo System (Biosemi, Amsterdam, The Netherlands) modified for passive recordings. Reference electrodes were placed on the shoulders of the subject.

### Signal processing

Following a previously validated signal processing pipeline (32), signals were band-pass filtered (1-60 cpm) and a Savitky-Golay filter (order 9, 1.7 s window) was applied. Slow-wave activation times were marked in the Gastrointestinal Electrical Mapping Suite (GEMS) (25) with subsequent manual review to ensure accuracy. Marked data files were exported for custom analysis of morphological feature collation and comparison, as described below.

### Morphology analysis

Averaged waveforms were extracted using an adaptive windowing technique. Adaptive windows were defined between some period preceding (*t*_pre_) and following (*t*_post_) each marked activation time, aligned to occur at *t* = 0. The values of *t*_pre_ and *t_post_* were calculated on a per-channel and per-subject basis using 20% and 80% of the mean slow-wave interval, respectively. After applying adaptive windowing, the median slow wave was calculated for each electrode with a detectable signal.

Downstroke amplitude, width, and recovery duration were calculated using validated algorithms for each windowed slow wave (24,32,33). We also introduced a new area-based metric to quantify waveform morphology. Specifically, the recovery stroke area (RSA) bounded by the normalized slow-wave signal (scaled 0 to 1), *s(t)*, and the baseline level, *b*, was integrated from ‘just prior’ the activation time (*t* = -*w*/2, half the downstroke width) until the end of the waveform *t* = *t_post_*.

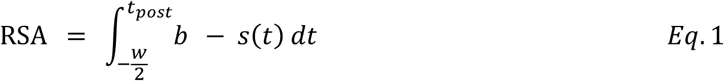

To account for differences in slow-wave periods, RSA was normalized according to Eq. 2:

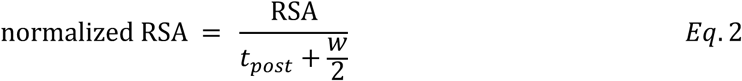

The morphological measures are summarized in Figure 6.

**Figure 6:**
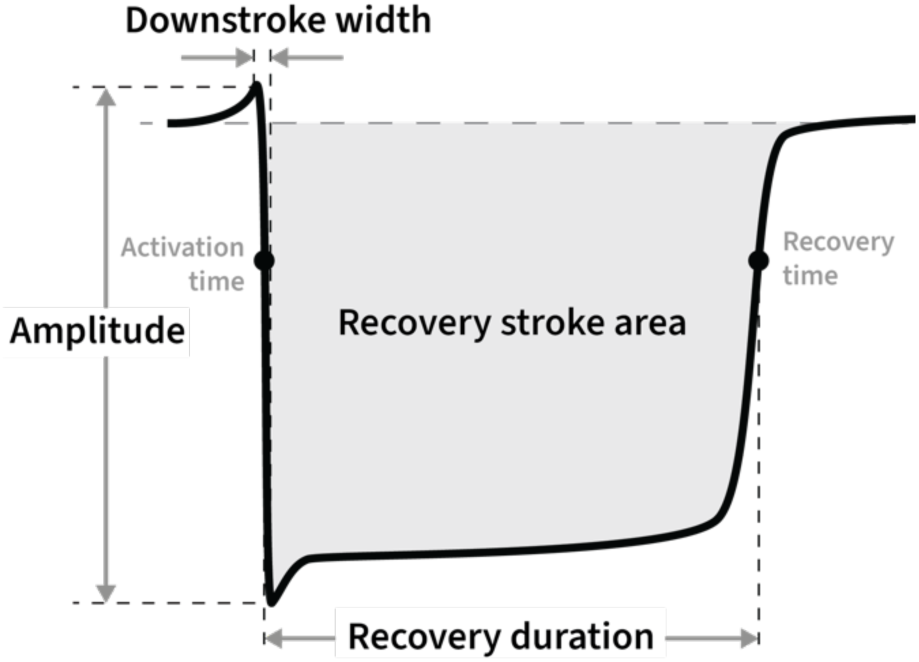
Slow-wave morphology measures. Amplitude was defined as the peak-to-peak voltage measure of the downstroke, downstroke width was the duration of the downstroke, recovery duration reflected the time between the activation and recovery times (defined as the maximum absolute gradient in the downstroke and recovery stroke, respectively), and recovery stroke area (RSA) was defined as the grey region indicated.

### Multiscale mathematical model

The effects of the underlying changes in slow waves contributing to the morphology of the extracellular potential were investigated using a previously described one-dimensional (1D) model (34). Briefly, the monodomain equation (Eq. 3) was applied to simulate the entrainment of gastric slow waves generated by the ICC network over a 345 mm line, representing the greater curvature of the human stomach. The initial 76 mm (22%) of the 1D model was modeled as the fundus, containing no slow-wave activity.

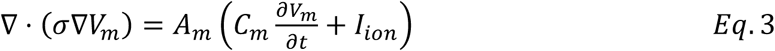

Where A*_m_* is the membrane area, C*_m_* is the membrane capacitance, σ is the conductance, V*_m_* is the membrane voltage, and I*_ion_* is the ionic current as estimated by a previously described cell model (34). Two gradients were prescribed to the 1D model in the antegrade direction. The first described the resting membrane potential gradient observed between the apex of the fundus and the pylorus (–56 to –70 mV (35)), and the second captured the intrinsic frequency of spontaneous depolarizations between the natural pacemaker region (76 mm distal of the apex of the fundus) and the pylorus (3 to 1.5 cpm (35)).

Simulations were performed for each cohort until solutions reached steady state. The resultant extracellular potentials were calculated based on the integration of membrane potentials across the gastric musculature at the midpoint of the 1D model (172.5 mm), as previously described (8,36). The parameters of the underlying cell model were adjusted until the calculated extracellular potentials reflected the morphology of each cohort.

To investigate the influence of morphology on global gastric function, the 1D model was extended to three dimensions, using an anatomically informed 3D human stomach geometry (37). The effects of the changes in morphology on gastric slow-wave propagation were compared to the baseline simulation once both models were solved until steady state.

### Statistical analysis

Linear mixed-effects models were fit to the data with cohort membership as a fixed effect, and subject and electrode number were assigned as random effects. Electrode number was nested within each subject. The normality of the model residuals was assessed by calculating skewness and visualizing the histogram and Q-Q plot. Homoscedasticity was assessed by visual inspection of the residual scatter plot. In cases where residuals were not normally distributed, data were square root transformed, and the linear mixed-effects model was refitted. Significance of cohort-wise differences was estimated using the R package *lmerTest* (38). Fixed effects were estimated using the R package *emmeans* (39) and are reported as estimated marginal means with 95% confidence intervals. A threshold of *p* < 0.05 was used for significance. All analyses were performed using R Statistical Software (v4.4.1) (66).

## CONCLUSION

This study presents a case for the value and potential functional influence of slow-wave morphology in gastric motility disorders. A new integral-based biomarker was introduced, along with a definition of typical morphological features to support the standardization of future slow-wave morphology research.

## Acknowledgements

The authors thank the clinical research and operating room staff at Auckland City Hospital, Auckland, New Zealand, and the University of Mississippi Medical Center, Jackson, MS, USA. The authors also thank Dr. Thomas Abell (now of University of Louisville), Dr. Christopher Lahr (retired), Mr. Simon Bull, Dr. Rachel Berry, Prof. Leo Cheng, and Prof. Greg O’Grady for their contributions to the original data collection. Finally, the authors would like to thank Dr. Armen Gharibans and Prof. Greg O’Grady for their critical review of the manuscript.

## Notes

**Funding Statement:** This study and/or authors were supported by a Rutherford Discovery Fellowship and Marsden Fund grant from the Royal Society Te Apārangi, New Zealand, the New Zealand National Science Challenge on High-Value Nutrition, the Health Research Council of New Zealand, and grants from the Ministry of Business, Innovation and Employment’s Catalyst: Strategic fund.

### Competing Interest Statement

No commercial or financial support was received for any material presented in this paper. TRA-G is a shareholder in FlexiMap Ltd. JCE and PD are shareholders in Alimetry (Auckland, New Zealand).

